# Genetic barcodes for ash (*Fraxinus*) species and generation of new wide hybrids

**DOI:** 10.1101/2024.02.19.581010

**Authors:** William J. Plumb, Laura J. Kelly, Joe Mullender, Robyn F. Powell, Laszlo Csiba, Miguel Nemesio-Gorriz, David Carey, Mary E. Mason, William Crowther, Jennifer Koch, Gerry C. Douglas, Richard J. A. Buggs

**Affiliations:** School of Biological and Behavioural Sciences, Queen Mary University of London, London, UK; Royal Botanic Gardens, Kew, Richmond upon Thames, UK; Forestry Development Department, Teagasc, Ashtown, Dublin, Republic of Ireland; United States Department of Agriculture, Forest Service, Northern Research Station, Delaware, OH, USA

## Abstract

Native ash tree species in Europe and North America are being devastated by ash dieback and the emerald ash borer, respectively. As worldwide ash species differ in their level of susceptibility to these threats, hybrid breeding may allow resistance to be transferred among species. However, we do not know the extent to which distantly related ash species can be crossed, and many ash species are difficult to identify from morphology alone leading to some mislabelling in living collections. Here, we develop a genetic barcode system for the identification of *Fraxinus* species based on three low-copy-number protein coding genes. We also conduct experimental crosses among ash species in different sections.

Our barcodes are effective in identifying ash samples to sectional level and in some cases to species level, and can also identify hybrids. They highlight that *F. mandshurica, F. platypoda* and *F. chiisanensis* may be frequently mistaken for one another in living collections. We succeeded in generating ten wide hybrid plants: two of section *Melioides* (species: *F. pennsylvanica*) □ section *Fraxinus* (species: *F. excelsior*) and eight of section *Ornus* (species unclear) □ section *Fraxinus* (species: *F. excelsior*). One hybrid from each of our crosses has survived natural infection with the ash dieback pathogen in Ireland. We also discovered a hybrid between section *Melioides* (species: *F. latifolia*) □ section *Fraxinus* (species: *F. excelsior*) formed spontaneously in the ash collection at Kew. Our findings facilitate the deployment of global ash species diversity in response to alien pests and pathogens.

**Societal impact statement:** The world-wide diversity of ash trees includes genetic information encoding resistance to the ash dieback fungus and the emerald ash borer beetle, which are currently devastating ash populations in Europe and North America. In order to mobilise this genetic diversity in conventional breeding programmes we need to be able to accurately identify ash species from around the world, and cross them with one another. Here, we present a genetic barcoding system for ash species, and a series of hybridisation experiments between European ash and other species. Two of the hybrids show early promise against ash dieback.

## Introduction

In the temperate regions of Europe, European ash (*Fraxinus excelsior*) is an ecologically and economically important broadleaf tree species (Dobrowolska *et al*., 2011; Mitchell *et al*., 2014) but is severely affected by the rapid spread of the non-native fungal species, *Hymenoscyphus fraxineus*, causing symptoms commonly referred to as “ash dieback” (ADB) or “Chalara” (Pautasso *et al*., 2013; Gross *et al*., 2014; Coker *et al*., 2019). European ash populations are further threatened by an insect species, the Emerald Ash Borer (EAB) (Orlova-Bienkowskaja & Bieńkowski, 2018) which has already devastated ash populations in North America (Herms & McCullough, 2014). Worldwide species of ash differ in their levels of resistance to ADB and EAB (Koch *et al*., 2007; Kowalski *et al*., 2015; Rigsby *et al*., 2016; Nielsen *et al*., 2016; Plumb *et al*., 2019; Kelly *et al*., 2020; Koch, 2025). Thus some non-native *Fraxinus* species could be suitable to replace susceptible populations in Europe and North America (Jepson & Arakelyan, 2017; Marzano *et al*., 2019; Lévesque *et al*., 2023) or used as parents in hybrid breeding programmes to transfer resistance among species (Koch *et al*., 2007, 2012; Plumb *et al*., 2019).

Several classifications of the genus *Fraxinus* have been published (e.g. Wenzig, 1883; Wesmael, 1892; Lingelsheim, 1907; Miller, 1955; Nikolaev, 1981; Jeandroz *et al*., 1997; Wallander, 2008, 2012; Hinsinger *et al*., 2013) with differing species divisions and sectional divisions. In this paper we follow the classification of Wallander (2012), where the genus contains over 40 species, many of which are placed into six sections. Our only exception to this is that we do not place *F. griffithii* into section *Ornus* due to its placement outside the *Ornus* clade in the phylogeny of Kelly *et al*. (2020) and Hinsinger *et al*. (2013); we treat it as *incertae sedis*. We note that Hinsinger et al (2013) places *F. platypoda*, *F. chiisanensis* and *F. cuspidata* in section *Melioides s. l.*. By contrast, in Jeandroz *et al*. (1997) *F. platypoda* is placed as a sister to *F. mandshurica* and *F. cuspidata* is in a polytomy with clades containing species of Wallander’s sections *Fraxinus* and *Melioides*. In Kim et al (2022), *F. chiisanensis* and samples of *F. platypoda* from Japan are placed in *Melioides s.l.* but samples of *F. platypoda* from China are placed in section *Fraxinus* where they are in a clade with *F. mandshurica*. In Wallander (2012) *F. platypoda*, *F. chiisanensis* and *F. cuspidata* are *incertae sedis*, and this is how we treat them here.

*Fraxinus* species can be difficult to tell apart as many of their most important morphological features are only visible at certain times of year, or are age-dependent (Henry, 1914; Wallander, 2012). Some closely-related species are morphologically similar and have been defined partly on the basis of their geographic distribution. Some species show high intraspecies variability and phenotypic plasticity. It is hard to differentiate between some *Fraxinus* species using DNA sequencing of the internal transcribed spacer (ITS) of nuclear ribosomal DNA, as sequence similarity is high between closely related species and intragenomic variation is present within some individuals (Wallander, 2008). Ploidy level (and hence C-value) is a useful method to distinguish among certain species but most species are diploid and others, such as *F. caroliniana*, include more than one ploidy level (Whittemore *et al*., 2018).

Hybridisation occurs between some species of *Fraxinus.* The sister species *F. excelsior* and *F. angustifolia* naturally hybridise in Europe (Fernández-Manjarrés *et al*., 2006; Heuertz *et al*., 2006; Thomasset *et al*., 2011) and have also been crossed by breeders (Raquin *et al*., 2002). Many of the *Fraxinus* species in the section *Melioides* clade hybridise in areas of coexistence in North America (Russell M. Burns, 1990; Wallander, 2008). There is anecdotal evidence of natural hybridisation in regions of species overlap between section *Melioides* and section *Ornus* (Wallander, 2012). Henry (1914) documented various artificial interspecific crosses of *Fraxinus* species, though the extent to which they were successful is unclear. Hybrids of *F. nigra* □ *F. mandshurica* (both within the section *Fraxinus*) were produced by Davidson and two commercial clones, *Northern treasure* and *Northern gem*, were patented (Davidson, 1999; Davidson & Ronald, 2001). Koch et al. have successfully produced new *F. nigra* □ *F. mandshurica* hybrids and second-generation backcrosses appear to be resistant to EAB (Koch *et al*., 2007, 2010). Within-section hybrids between *F. mandshurica* and *F. angustifolia* subsp. *syriaca* (syn. *F. sogdiana* used by authors of the study) have been produced and propagated using embryo culture, with one clone showing photosynthetic heterosis (He *et al*., 2021, 2023). Wright (1953) produced hybrids between *F. pennsylvanica* and *F. velutina* (both in section *Melioides*). A cross between *F. velutina* and *F. pennsylvanica* has been used to map candidate loci for salt tolerance (Liu *et al*., 2024). Wider hybrids between sections *Fraxinus* and *Melioides* have also been produced. Johnson and Heimburger (1946) reported hybrids between *F. excelsior* var. *aureovariegata* (section *Fraxinus*) and *F. americana* (section *Melioides*), *F. pennsylvanica* (section *Melioides*) and *F. quadrangulata* (section *Dipetala*). Hybrids have been produced in China involving *F. mandshurica* and *F. americana*, and *F. mandshurica* and *F. velutina*, using electrostatic treatments to overcome fertility barriers; these hybrids showed heterosis in growth (Zeng *et al*., 2015; He *et al*., 2019; Cao *et al*., 2023).

Hybrid breeding programmes have been set up in response to other well-established pathogens of trees, such as Dutch elm disease (Smalley & Guries, 1993; Santini *et al*., 2007) and chestnut blight (Anagnostakis, 2012), producing trees with increased resistance. The process of interspecific hybridisation is usually not as simple as identifying two trees of different species and cross-pollinating them. Prezygotic incompatibilities include: flowering time differences between the species, failure of pollen germination on the stigma of a different species, and reduction or halting of pollen tube growth. Post-zygotically, the hybrid endosperm or embryos may fail to develop. Interventions may allow some of these barriers to be overcome, including: pollen storage, mentor pollen (Knox *et al*., 1987) and embryo rescue (Sharma *et al*., 1996; Raquin *et al*., 2002).

Whilst considerable effort may be invested in producing novel hybrids among tree species, hybrids may form spontaneously in arboreta and botanic gardens (M Maunder, C Hughes, JA Hawkins, A Culham, 2004). Such hybrids have often been viewed as something to be avoided (Ensslin & Godefroid, 2019), especially if they are hard to identify. However, if we can easily discover hybrids and identify their parental species using molecular markers, such spontaneous hybridisation events could become a valuable resource in plant breeding.

The growing interest in research programmes that involve different species of ash and their hybrids (Koch *et al*., 2007, 2012; Zeng *et al*., 2015; Plumb *et al*., 2019; Kelly *et al*., 2020; He *et al*., 2021; Cao *et al*., 2023; Liu *et al*., 2024) means that a method for their easy identification, usable on any tissue type at any time of year, is highly desirable. Whole-genome sequences for 22 *Fraxinus* species have recently been published (Kelly *et al*., 2020). This provided us with the opportunity to undertake genome-wide searches for nuclear loci suitable for barcoding within the genus. Here, we describe development of a set of barcoding regions using single-copy genes in *Fraxinus*. We show that these barcodes can identify specimens to sectional level and often to species level. We also produce new inter-sectional *Fraxinus* hybrids, and verify their parentage using our new barcodes.

## Materials and Methods

### *Fraxinus* accessions

We collected materials from *Fraxinus* accessions in arboreta and botanic gardens (Supplementary Table 1) and from the United Stated Department for Agriculture Forest Service ash breeding program at the Northern Research Station (in Delaware, Ohio). We focused on these, rather than herbarium specimens, as they are potential sources for: experiments, future planting stocks and hybridisation programmes. Some of these specimens were used in separate studies as sources of scion material to set up assays for EAB susceptibility in Ohio (Kelly *et al*., 2020) and trials for ADB susceptibility in England (see: www.forestresearch.gov.uk/research/testing-a-range-of-ash-species-for-tolerance-to-ash-dieback/). The living collections in the British Isles that contributed materials were: Royal Botanic Gardens Kew (including Wakehurst), Cambridge University Botanic Garden, Westonbirt Arboretum, Earth Trust, RHS Wisley, Ness Botanic Gardens, Royal Botanic Garden Edinburgh, Dawyck Botanic Garden, The National Botanic Gardens Glasnevin (Ireland), The John F. Kennedy Arboretum (Ireland) and a private collection belonging to Gerry Douglas (Dublin, Ireland). We purchased from Sandeman Seeds (Dalkey, Ireland) seeds labelled as *F. mandshurica*, and germinated them in Ireland. We did not have access to some accepted taxa of *Fraxinus*, mainly due to poor representation of species native to Mexico and South America in British and Irish collections. As these species do not grow well in European conditions, they are unlikely to be part of the solution to ADB and EAB. DNA was extracted from the leaves or cambial tissue of specimens using Qiagen DNeasy® Plant Mini Kit (QIAGEN Ltd., Manchester, UK) or CTAB protocols (Kelly et al., 2020).

### Selection of candidate barcoding genes

A genome assembly for *F. excelsior* was published in 2017 (Sollars *et al*., 2017) based on a tree grown from seed collected in a woodland in Gloucestershire, UK. For a further 28 of the samples we used, Kelly et al (2020) had previously sequenced whole genomic DNA (see Column E in Supplementary Table 1) and produced genome assemblies. The species identifications of these samples had been checked by Dr Eva Wallander (an expert on the genus) based on morphology and ITS, and where these differed from the labels in the living collections we took the identification by Wallander to be correct. In the case of accessions 1973-6204 and chi-12 there was uncertainty about the identity and Kelly et al (2020) initially labelled them as *Fraxinus* sp. From the whole genome assemblies of these 29 *Fraxinus* samples, Kelly et al (2020) had identified groups of putatively orthologous protein coding gene sequences in them and three outgroups (*Olea europaea*, *Erythranthe guttata* and *Solanum lycopersicum*) using OMA (Altenhoff *et al*., 2019). Full details of methods can be found in Kelly *et al*. (2020), but briefly, the full-length sequences (i.e. including both exons and introns, where present) for OMA groups that included sequences from all 29 *Fraxinus* samples and the three outgroups were aligned using MUSCLE (Edgar, 2004) via GUIDANCE (Sela *et al*., 2015) and unreliably aligned positions removed. OMA groups with alignments shorter than 300 characters in length, or which included sequences with <10% non-gap characters were excluded, leaving a set of 1385 OMA groups.

The alignments for these groups were examined using a pre-release version of CONTEXT, a phylogenomic dataset browser (v0.8. pre-release) (https://github.com/lonelyjoeparker/qmul-genome-convergence-pipeline/blob/master/CONTEXT.md) to generate statistical information about each OMA group including longest sequence length, longest un-gapped region, number of variant nucleotides and amino acids, and the mean entropy values for nucleotides and amino acids. We selected candidates for further analysis based on their: degree of variance (25% or above), greatest overall length, and length of the longest un-gapped region (being longer than 800bp). For these candidate OMA groups, maximum likelihood trees were generated using RAXML-ng version 0.8.0 (Kozlov *et al*., 2019), using the following parameter settings: --model GTR+G --tree pars{10},rand{10} --bs-trees 1000. We examined the trees for congruence with the species-tree from Kelly *et al*. (2020). We checked for putative paralogs by searching the sequence from *F. excelsior* against its reference genome (assembly version BATG0.5) using BLAST; OMA groups were excluded for if the sequence matched another region of the genome with total coverage of greater than 75% of the query sequence length (suggesting that one or more paralogs of the query gene was present in the genome). The sequences for the three outgroups were removed from the remaining candidate OMA groups and new alignments generated using the MUSCLE tool from EMBL-EBI (https://www.ebi.ac.uk/Tools/msa/muscle/ accessed October 2017).

### Primer design and testing

We selected four variable OMA groups for primer design. These were OG8679 (a pentatricopeptide repeat, according to the functional annotation of BATG0.5), OG9972 (a histidine phosphotransferase protein), OG22833 (a protein of unknown function) and OG8143 (a polyadenylate-binding protein 1). Primers were designed using Primer3web v4.0.0 (http://bioinfo.ut.ee/primer3/ accessed September 2017) and purchased from Eurofins MWG Operon (Ebersberg, Germany). The primers were provided lyophilised, and Milli-Q® H_2_O was used to reconstitute them to a concentration of 100 pmoles/μl. We tested these by performing PCR on our DNA extractions from 151 trees in living collections.

Each PCR reaction contained <100/ng genomic DNA, 1μl forward primer 10pmol/µl, 1μl reverse primer 10pmol/µl, 25μl DreamTaq Green PCR Master Mix (2X) (DreamTaq DNA Polymerase), 2X DreamTaq Green buffer, dNTPs, and four mM MgCl_2_ (Thermo Fisher), 10μl 5X TBT-PAR (containing trehalose, bovine serum albumin (BSA), and polysorbate-20 (Tween-20), and 2μl DMSO (Dimethyl sulfoxide). Reactions were made up to 50μl with Milli-Q® H_2_O. The conditions used were: an initial denaturation stage at 94°C for 6 min; followed by 38 cycles of denaturation at 94°C for 1 min, annealing at 52°C for 1 min and extension at 72°C for 2 min; with a final extension at 72°C for 7 min. PCRs were carried out on a GeneAmp® PCR System 9700 thermocycler (Applied Biosystems, Warrington, UK).

Agarose gel electrophoresis was used to determine the presence of correctly sized PCR products. Gels were prepared by dissolving 1.5g molecular grade agarose (Bioline Reagents Ltd., London, UK) in 150ml 0.5x Tris-borate EDTA (TBE) (27 g of Tris base, 13.75 g of boric acid and 10 ml of 0.5 M Ethylenediaminetetraacetic acid (EDTA) (pH 8.0), made up to 1 litre using Milli-Q® H_2_O) using a microwave to heat the solution. Once the agarose was fully dissolved it was left to cool to 55°C before mixing 6μl 10mg/ml ethidium bromide (EtBr) into the solution. The solution was then poured into a prepared gel tray and left to cool at room temperature until it had solidified. Once the gel had set, it was placed into a Mini-Sub® Cell gel tank (Bio-Rad Laboratories Ltd., Hemel Hempstead, UK) and submerged in 0.5x TBE buffer. Gels were run at 100V / 600mA for 30 min with the amplicon samples and a 1 Kb Plus DNA Ladder (Thermo Fisher). The gels were visualised using a UV trans-illuminator Gel Doc 2000 and Quantity One 1-D analysis v4.6.5 software (Bio-Rad).

For sequencing, the PCR products were purified using a NucleoSpin® Gel and PCR Clean-up kit (Macherey-Nagel). Sequencing reactions were made up of 20-25/ng of the purified PCR product, 0.5ul at 5pmol/µl of appropriate primer, 1ul 5x sequencing buffer, 0.25ul DMSO (Dimethyl sulfoxide) and 0.25ul BigDye® Direct Sanger Sequencing Kit Thermo Fisher Scientific UK Ltd (Loughborough). Cycle sequencing was performed using the following parameters: 35 cycles of denaturation at 96°C for 10 s, annealing at 52°C for 5 s and extension at 60°C for 4 min using a GeneAmp® PCR System 9700 thermocycler (Applied Biosystems, Warrington, UK). Samples were Sanger-sequenced using a 3730 Series Genetic Analyzer Thermo Fisher Scientific UK Ltd (Loughborough). The electropherograms were trimmed and the forward and reverse sequence for each sample were assembled into contigs using Geneious® ver8.1.9 software (Biomatters, Ltd., Auckland, New Zealand).

We aligned sequence data for the four barcode regions from all successfully sequenced samples using MUSCLE (Edgar, 2004) within JALVIEW (Procter *et al*., 2021). We trimmed the alignment to the ends of the outermost PCR primers. Each sample was given a label that began with an abbreviation of its taxonomic section (DIP = *Dipetala*, FRA = *Fraxinus*, MEL = *Melioides*, ORN = *Ornus*, PAU = *Pauciflorae*, SCI = *Sciadanthus*, INC = *Incertae sedis*), followed by its species ID, followed by its sequencing tube number. For the samples that had been used for whole genome sequencing and barcode primer design, the label contained the section and species allocated after identification by Eva Wallander using morphology and ITS data. For the other samples, the label contained section and species according to labels in living collections. We manually sorted the alignment by section and species. When this was done it was apparent that in all barcodes there were single nucleotide variant (SNV) alleles specific to sections and some species, but that some samples were anomalous. The anomalous samples were given the prefix ANOM and were reassigned to different sections according to the alleles they contained, with their new section abbreviation placed at the start of their label. Within sections, we then sought to manually cluster species, where possible, according to species-specific SNV alleles. Where a species included no samples that had been whole genome sequenced, we assumed that the majority of living collection labels that matched the correct section were also correct at a species level, allowing us to provisionally assign some of the anomalous samples to species. For each sample, we noted the SNV loci in each barcode that allowed assignment to section and, where relevant, species. Once we had as far as possible identified each sample (that had not been included in the whole genome sequencing panel) using barcodes, we compared these identifications, where possible, to independent identifications of the samples made by Eva Wallander using morphology and ITS, and also flow cytometry assessment of genome size.

### New hybrid crosses

In order to produce new interspecies crosses, we took scion material from a collection of mature *F. excelsior* plus trees selected from Irish forests for good forestry traits. These were: five predominantly female trees (Fex 32, 076, 8X and G 40) and five predominantly males (Fex 359, 42, 405-14 and G1). We took scion material from the John F. Kennedy Arboretum, Co. Wexford, Ireland, from trees labelled as: *F. pennsylvanica* (JFK.02931), *F. paxiana* (JFK.02928)*, F. floribunda* (JFK.02916), *F. japonica* (a synonym for *F. chinensis* in (Wallander, 2012))(JFK.02920)*, F. chinensis* (JFK.02906), *F. chinensis* subsp. *rhynchophylla* (JFK.02908), *F. retusa henryana* (a synonym for *F. floribunda* in (Wallander, 2012))(JFK2005.0132), *Fraxinus nigra* Marsh. × *Fraxinus mandshurica* (Canadian Hybrid No 8921 “Northern Treasure” JFK.2001.0159), *F. american*a (JFK.02899). From the National Botanical Gardens of Ireland, Glasnevin, we took scions from: *F texensis* (a synonym for *F. albicans* in (Wallander, 2012))(XX.011046.A2) and *F. mandshurica* (1934.011053.A2). We also took scion material from trees labelled as *F. mandshurica* (T1-T3) in Dr Gerry Douglas’ private garden, which were originally sourced from National Institute of Forest Science in Seoul, South Korea.

Scions were cut to length 15-20 cm and ∼0.5cm diameter, with at least two sets of lateral buds and one terminal bud. These scions were cleft-grafted to rootstocks at Teagasc, Kinsealy, Ireland between 2011- 2015 as follows. Two year old trees from naturally occurring Irish *F. excelsior* populations were sourced from a commercial nursery. During the period February-March, as the scion and rootstocks lay dormant, the rootstock trees were cut to 5-10 cm above soil level and an incision made to receive the scion. The basal end of the scion was cut to two opposing smooth-tapered cuts approximately 2-4 cm in length from the lowest set of buds using a sharp grafting knife. The cut scions were inserted into the rootstock cleft, with the cambium of the scions positioned into direct contact with the cambium of the rootstock. The graft union was bound using elastic grafting bands. The entire scion and graft union was dipped into molten paraffin wax (45°C). Grafted plants were potted into 3L pots with commercial-grade peat compost and maintained in the greenhouse. Sprouts that arose from below the graft union were routinely removed as these can lead to graft rejection. Grafted trees were later moved to the Teagasc Ashtown Research Centre, Ireland in 2018, following the Kinsealy site’s closure.

Cross-pollination was carried out between March and April of 2015 as some of the grafted trees came into flower. The grafted plants were at least two years old at the time of flowering and crossing. The following non-native accessions were used in the crosses: trees labelled as *F. mandshurica* in Dr Gerry Douglas’ garden were used as male parents (no female flowers were available), *F. pennsylvanica* (JFK.02931) used as a female parent, *F. chinensis* (JFK.02906) was used as a male and a female parent, *F. retusa henryana* (JFK2005.0132) was used as a female parent, *F. japonica* (JFK.02920) was used as a male parent, and *Fraxinus nigra* Marsh. □ *Fraxinus mandshurica* (Canadian Hybrid No 8921 “Northern Treasure” JFK.2001.0159) was used as a female parent. Each cross was with *F. excelsior* as the other parent. In addition to the grafted *F. excelsior* plus trees 076, 8X, G40, 32, 42, 405-14, G1, and 359, we also used *F. excelsior* 98, a selected plus tree which was growing outside nearby.

Pollen was collected from male flowers at anthesis by gently tapping inflorescences so that pollen fell into glass petri dishes (60 x 17mm; Thermo Fisher). Dishes were labelled with the pollen donors’ accession number. Glass was used instead of plastic petri dishes for collection and storage, to avoid static electricity making the pollen challenging to manipulate. Pollen was stored for future use by wrapping parafilm around the petri dish’s lid, leaving a small opening for drying. Wrapped dishes were then placed in a glass desiccator containing silica gel and stored in a refrigerator at 4°C until required in the same season. In some cases when the non-native species were hermaphrodite, the male flowers were emasculated to prevent self-fertilisation. Pollinations were performed at least twice for each cross using pollen collected on the same day or from refrigerator-stored pollen and within a few days of the initial pollination event. Stigmas on selected female trees were checked daily over the period in which the inflorescence was maturing and pollinated when some of them showed a glistening surface. Pollen was applied using a tiny paintbrush that was lightly brushed against the stigmas to release a puff of pollen. A tag was placed around each inflorescence of the pollinated tree, with an allocated number corresponding to the parental details for the cross. In one case two different pollen donors *F. excelsior* 405-14 and *F. excelsior* G1 were used. Pollen of *F. excelsior* 405-14 was used on the first day and pollen of *F. excelsior* G1 was used on a subsequent day for the same cross as ‘mentor pollen’ may facilitate hybridisation in some cases. The pollination period was generally in the last two weeks of March when stigmas were deemed to be most receptive. After the first pollination event the trees were moved to a separate section of the greenhouse to minimise potential contamination by nearby ash trees which may have been flowering. After that, the pollinated trees were grown in the glasshouse for the summer period in which unpollinated pistils abscised naturally. As *F. excelsior* plus tree 98 was field grown, the pollinated female inflorescences were enclosed and sealed in pollination bags and unbagged inflorescences were used as open pollinated (OP) intra-species controls under the assumption that any ambient pollen would be from *F. excelsior*.

The pollinated inflorescences were observed for the development of the samaras over several months during which many immature/unfertilised samaras abscised naturally. During the last week of August and the first two in September, an assessment was made of seed production from crosses. Full samaras were excised, and surface sterilised in 7% (w/v) calcium hypochlorite Ca(ClO)_2_ solution for 20 min, followed by three rinses in sterile water and aseptic air drying. Thereafter the samaras were dissected aseptically under a stereomicroscope, and the seeds were removed and distributed onto B5 medium for germination (Gamborg et al. 1968) in small Petri plates (5 cm) containing 6-7 ml of medium with sucrose in concentrations of 2%, 3%, 4%, 5% (w/v) and solidified with phytagel 3 g L^-1^. Culture conditions were at 22 ± 1°C with 16-hour photoperiod using Philips® cool white fluorescent tubes giving a photon flux density of 58.6 µmol m-1s-1. Seed germination and viability were assessed after approximately six weeks and after this period, ungerminated seeds were either transferred to fresh media, or the embryos were excised from seeds for embryo rescue culturing. In some cases we did not attempt whole seed germination but embryos were excised and cultured directly after seed collection and surface sterilisation. Embryo rescue was performed under a stereomicroscope in a laminar flow hood under aseptic conditions. An incision was made along the longitudinal axis of the seed and into the endosperm and the outer seed coat was peeled off. The seed was prised open at the cotyledon end to reveal the embryo attached within one half of the seed. The embryo was gently prised off the endosperm using the tip of a scalpel and placed directly into B5 medium. Whole seeds and isolated embryos were transferred to fresh media every 6-8 weeks at least six times. We routinely used B5 medium with 3% sucrose in the latter culture periods. Germinated seedlings were transferred to glass jars (volume, 150 ml) on WPM medium salts (Lloyd and McCown 1980) and vitamins of B5 medium (Gamborg et al. 1968) supplemented with 3% w/v activated charcoal and then weaned to the glasshouse.

In 2018, leaf material was collected from each hybrid for DNA extraction. The *Fraxinus* genetic barcodes were used to identify successful new hybridisations. DNA extraction, PCR and sequencing was performed in the same way as it was for the living collections (see above). Sanger sequence reads were then trimmed, cleaned and assembled into contigs. The sequence chromatograms were viewed using Geneious prime version 8.1 and we looked for double peaks (i.e. heterozygosity) at SNP loci that differed between the putative parental species sequenced in the test panel. We carried out an initial screening using the barcode region OG9927 and hybrids found with this locus were also sequenced with the barcoding regions OG22833 and OG8143. From 2018 onwards the new hybrids were grown in an area of high ADB disease pressure at the Teagasc Ashtown Research Centre, Ireland, and records were kept of death of trees due to ADB.

### Seeking spontaneous hybrids

As hybrids may spontaneously form in living collections, we collected four volunteer seedlings growing within the main ash collection area at Kew Gardens, and used the barcodes to test if they were hybrids.

### Ploidy investigations

Where polyploidy has been reported for certain *Fraxinus* taxa, measurement of genome sizes using flow cytometry can aid in the identification of species and their hybrids. In such cases, nuclear DNA contents were estimated following the one-step flow cytometry procedure (Dolezel *et al*., 2007).

Approximately 1 cm^2^ of the sample leaf material was incubated for 30 seconds on ice in 1 ml of ‘general-purpose buffer’ (GPB) (Loureiro *et al*., 2007) supplemented with 3% PVP-40, after which leaf material from the calibration standard *Petroselinum crispum* or *Pisum sativum* was added (Obermayer *et al*., 2002). This material was chopped together rapidly using a fresh razor blade. Another 1 ml of the isolation buffer was then added to the chopped material, and the homogenate was filtered through a 30µm nylon mesh (Celltrics 30µM mesh, Sysmex, Goritz, Germany). Following filtering, 100 μl propidium iodide (1 mg/mL) was added, and the sample was incubated on ice for 10 minutes before analysis. The relative fluorescence of 5000 particles was then recorded using a Partec Cyflow SL3 flow cytometer (Partec GmbH, Münster, Germany) fitted with a 100 mW green solid-state laser (532 nm, Cobolt Samba, Solna, Sweden) and the output histograms were analysed with the FlowMax software v.2.4 (Partec GmbH, Münster, Germany).

## Results

### Testing of barcodes

At least one barcode region (Table 1) amplified successfully and yielded sequence data for 148 DNA samples from 45 *Fraxinus* species or subspecies (see Supplementary Table 2). Usable sequences were generated for 143 samples with OG9927, 135 samples with OG22833, 133 samples with OG8143 and 132 for OG8679. If DNA quality was poor, the second primer sets for OG9927 and OG8679 tended to produce lower-quality reads. Trimmed alignments of all clean sequences generated can be found in Supplementary Files 1-4, ordered by section and species. These are also available at https://doi.org/10.5281/zenodo.10066755. Examination of these alignments showed that all of the barcodes contained SNV alleles that were specific to taxonomic sections and present in all samples within those sections. There were also many SNV alleles that were specific to species, though these were not always present in all samples from a species.

**Table 1.**
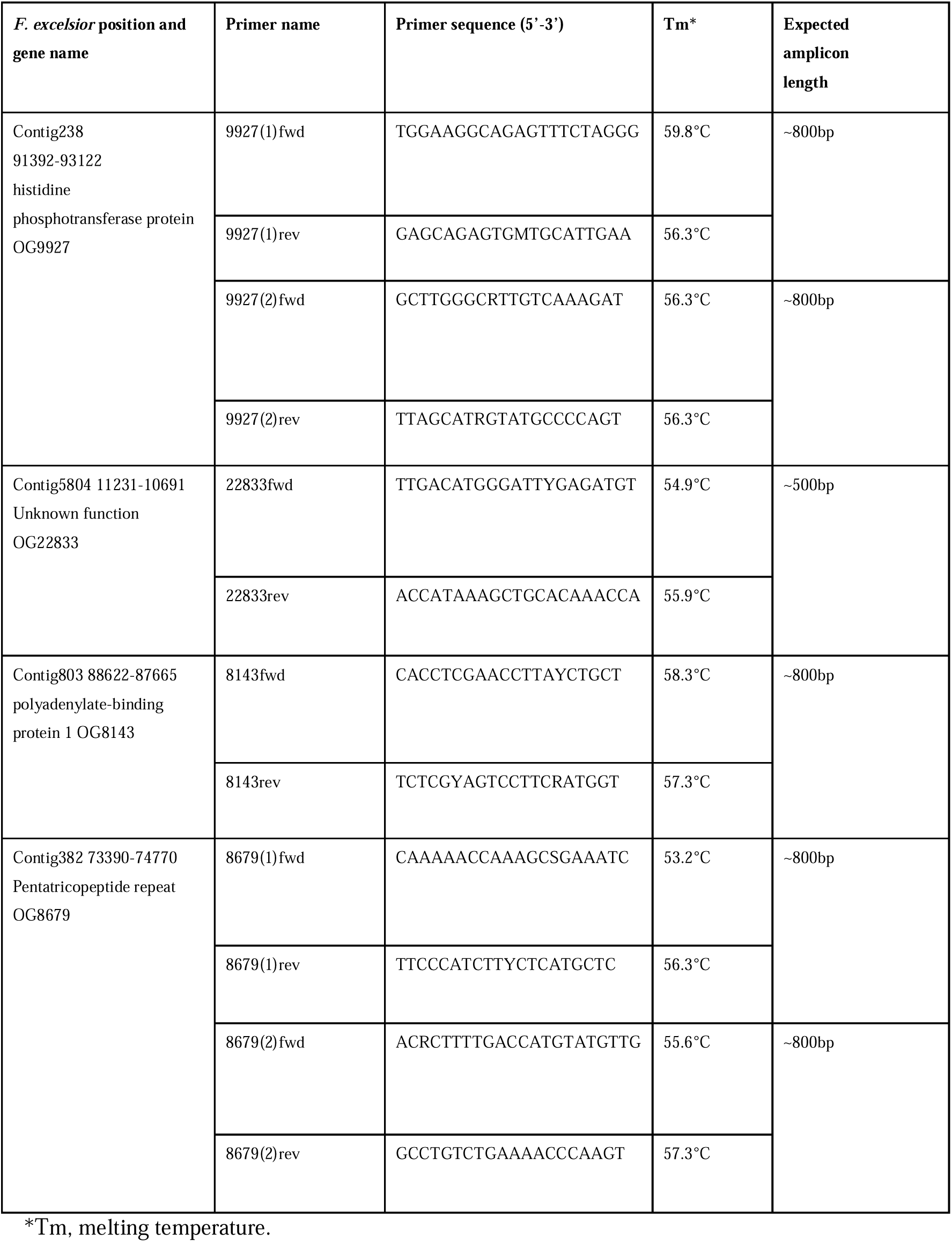
Primers designed and tested for *Fraxinus*species barcoding. Each primer’s location is given based on the *F. excelsior* BATG0.5 genome. OG numbers refer to the Ortholog group number assigned to genes in Kelly et al 2020.

Supplementary Table 1 gives a list of all tree accessions used, and details of the extent to which they can be identified to section and species levels using each DNA barcode. Barcodes OG9972, OG22833 and OG8143 gave conflicting sectional results for only two samples. In contrast, barcode OG8679 gave many double peaks in chromatograms and gave sequence results that sometimes conflicted with the other barcodes for nine samples. In the sequence alignment for OG8679, several SNP loci had three or even four alleles present among the samples. To give one example, site 546 was: G in the majority of samples, T in all four samples of *F. paxiana*, one of eight individuals of *F. floribunda*, one of six individuals of *F. excelsior,* C in two of three individuals of *F. lanuginosa*, A in all individuals later identified as *F. chiisanensis* (except one which was R (A/G)) and one of eight individuals of *F. quadrangulata* and in two of four individuals of *F. platypoda*, R in one of seven *F. nigra* and one of five *F. latifolia*. Those individuals with an A at position 546 all had an A at position 592, a G at position 657 and a T at position 674. Those that had an R at position 546 had an A or R at position 592, a K at position 657 and a T or Y at position 674. Such a pattern is best explained by the presence of two paralogs, with some samples showing amplification of both, and some only amplifying one copy, potentially due to loss or mutation of the primer site at the other copy. There were no cases in which OG8679 was able to give us sectional or species assignments for a sample that were unavailable from one of the other barcodes. Therefore we decided to discard OG8679 as a barcode and focus only on OG9972, OG22833 and OG8143.

Using OG9972, OG22833 and OG8143, 148 samples could be assigned to section (taking a majority vote in the three cases of sectional conflict) and 88 could be assigned to species. Those 60 samples that could not be identified to species level were all in the sections *Melioides*, *Fraxinus*, *Dipetalae* or *Ornus*. These sections are known to have similar species that sometimes hybridise. No diagnostic alleles were found for *F. excelsior* (in section *Fraxinus*), the most widespread species in Europe, or *F. pennsylvanica* and *F. americana* (in section *Melioides*), the most widespread species in North America, probably due to widespread hybridisation with other species found within their ranges.

Of the 148 samples assigned to section or species level by the three barcodes, 67 were also identified by Eva Wallander using morphology and ITS. Of these, none were assigned to a different section by the two methods of identification. For 34 samples we gained species level identifications using both methods. In 38 cases these agreed. The cases of disagreement were for *F. pennsylvanica*/*F. caroliniana*/*F. velutina* which are all closely related within section *Melioides*, or *F. chinensis*/*F. baroniana*/*F. floribunda* in section *Ornus*.

All three barcodes were each able to identify most of the 148 samples to sectional level. However, section *Fraxinus* was hard to identify using OG22833 and OG8143. For section *Fraxinus,* OG9972 performed best, with three clear section-specific alleles (Table 2). Overall, OG9972 identified 54 samples to species level, OG22833 identified 65 samples to species level, and OG8143 identified 47 samples to species level. Within section *Dipetala*, OG9972 performed best at identifying species and OG8143 could not distinguish any species. Within section *Fraxinus*, OG9972 and OG22833 both performed better than OG8143, but OG8143 was particularly useful for distinguishing *F. mandshurica*. In section *Melioides*, all barcodes were unable to distinguish species, with the exception of *F. velutina*, which could be identified by OG22833. In section *Ornus*, the barcodes differed widely in their patterns of diversity: OG9972 was good for identifying *F. floribunda* and *F lanuginosa*; OG22833 was good for distinguishing *F. chinensis* and OG8143 was good for distinguishing *F. chinensis* and *F. ornus*. Within the small section *Pauciflorae*, all barcodes could distinguish *F. greggii* but only OG8143 could distinguish *F. gooddingii*. All barcodes performed well at distinguishing the species not assigned to sections (*Incertae sedis*) but OG8143 was alone in being unable to identify *F. griffithii* to species level.

**Table 2.**
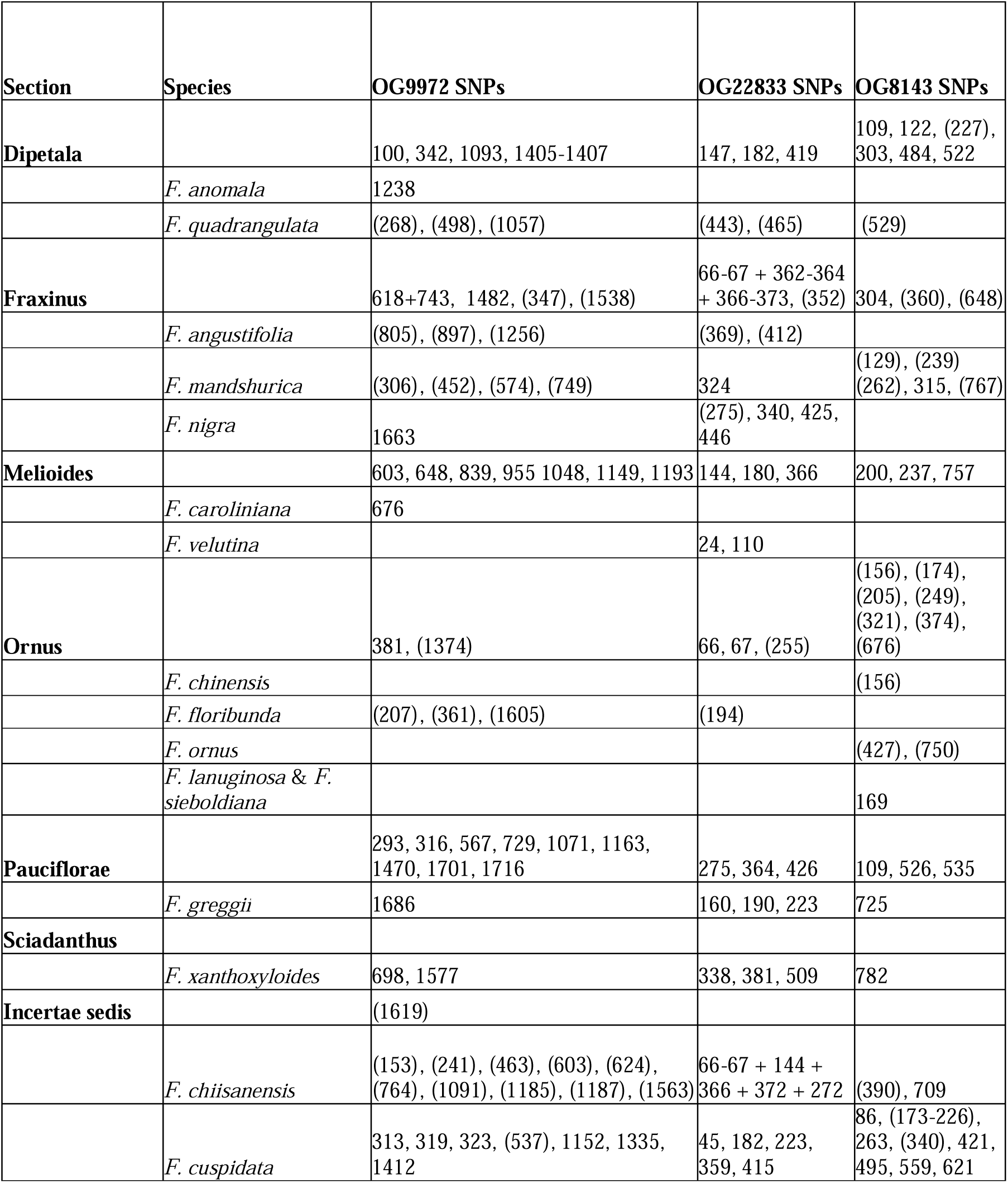
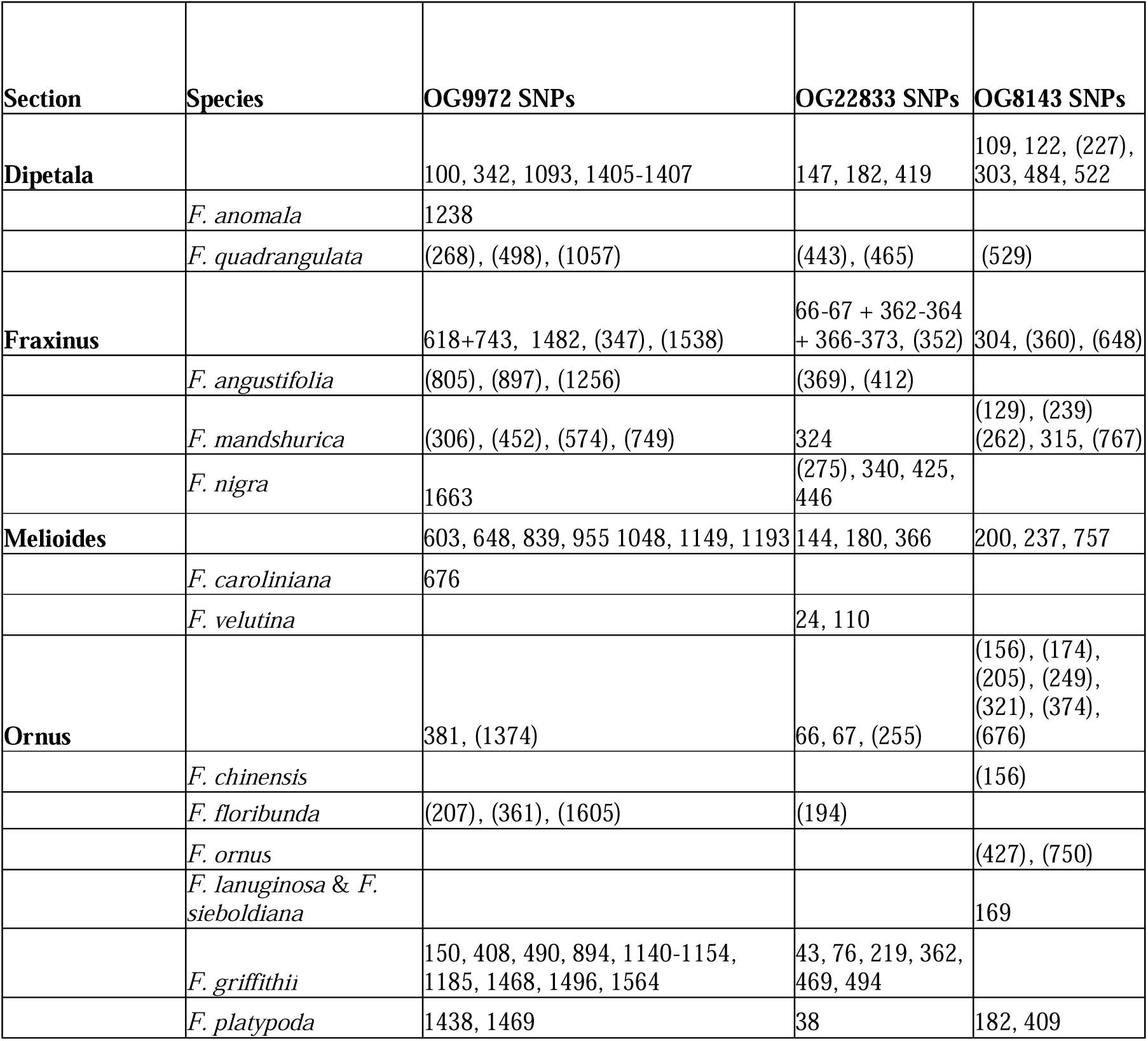
Diagnostic SNV allele locations for sections and species of the genus *Fraxinus* using three barcodes. Locations refer to positions in the alignments in Supplementary Files 1-3, where the first position in the first base in the forward primer. Locations in brackets mean that there is an allele unique to the species but not fixed within the species, so its presence is diagnostic but its absence is not. The plus sign is used for loci that must be in a particular combination to identify a species or section. The sections are according to Wallander (2013) classification with the exception of *F. griffithii* which we place as *Incertae Sedis* due to its placement in the species-tree inferred in Kelly *et al*. (2020).

Of our 148 sample identifications with barcodes, 25 conflicted with the labels in the living collections that they were derived from. Nineteen of these conflicts involved assignments of different sections, and six were assignments of different species within sections. Of the 19 sectional conflicts, four identifications were also available from Eva Wallander who gave sectional classifications that agreed with the barcodes.

There were two cases where the tree label said *F. platypoda* (*Incertae sedis*) but the barcodes indicated *F. mandshurica* (Fraxinus). These were JFK.02932 and USDA FS plat-3, and for the latter, the barcode identification was confirmed by Wallander. The barcodes assigned Kew 1900-44501 *F. quadrangulata* (*Dipetala*) and Kew 2011-1622 *F. quadrangulata* (*Dipetala*) to section *Fraxinus*. A sample labelled *F. greggii* (Cambridge 19860253*A) was identified as *F. cuspidata* (*Incertae sedis*) by the barcodes. A sample labelled *F. chinensis* subsp. *chinensis* (*Ornus*)(Westonbirt 56.0455) was identified as *F. pennsylvanica* (*Melioides*) by both the barcodes and Wallander. A sample labelled *F. platypoda* (*Incertae sedis*)(USDA FS plat-1) was identified as *F. pennsylvanica* (*Melioides*) by both the barcodes and Wallander (it is now relabelled in the USDA FS collection as pe-103). Three samples labelled as *F. nigra* (*Fraxinus*)(Wakehurst 1996-5167 and 1996-5169 in replicate) were identified as section *Melioides* by the barcodes, and 1996-5169 was identified as *F. pennsylvanica* (*Melioides*) by Wallander. One sample labelled *F. americana* (Melioides)(JFK.02899) was identified as *F. chinensis* (Ornus) by the barcodes; morphological evaluation of this accession is needed. Seven samples labelled as *F. mandshuric*a (GD-T3, GD-T1 and Kew 1989-3691, 1989-8285, 1992-375, 1992-376 and 1992-377) in section *Fraxinus* were identified by the barcodes as being not in the section *Fraxinus*. Closer examination of their genome size and bud morphology (see details below) allowed them to be identified as *F. chiisanensis* (*Incertae sedis*). Five seedlings germinated in Ireland from seeds purchased from Sandeman seeds (Dalkey, Ireland) labelled as *F. mandshurica* were confirmed to be *F. mandshurica*.

Of the six cases where the barcodes assigned a different species within the same section as the living collection label, three of them were also examined by Wallander who in two cases made the same identification as the label (USDA FS PE_48 FRAX10 *F. pennsylvanica* which the barcodes assigned to *F. velutina* and USDA FS Bar-2 FRAX28 *F. baroniana* which the barcodes assigned to *F. chinensis*). In one case Wallander made the same identification as the barcodes (Kew 1973-6204 FRAX29 *F. bungeana* which the barcodes assigned to *F. ornus*), suggesting that the barcoding was accurate. The other three cases involved: JFK.02906 *F. chinensis* being assigned to *F. ornus*; Kew 2004-1231 *F. paxiana* and USDA FS BUN-5 *F. bungeana* being assigned to *F. chinensis*. Details of the SNP loci used for each assignment can be found in Supplementary Table 2 and seen in Supplementary Files 1-3.

*Fraxinus excelsior* (USDA FS ex-12) had several positions in OG9927 that were diagnostic for *F. cuspidata* (Incertae sedis) or were heterozygous for a base found in section *Fraxinus* and a base found in at least one individual of *F. cuspidata*, suggesting it could be a hybrid between these two species. However in OG22833 and OG8143 there was no evidence of this.

### Genome size measurements

Genome sizes were measured for several *Fraxinus* accessions. These results are shown in Supplementary Table 1, column Y, with inferred ploidy level from these genome sizes in column Z.

The majority of samples had 1C genome sizes between 700Mb and 1000Mb and were inferred to be diploid. Some, but not all, accessions identified by barcodes as *F. chinensis* were hexaploid: Westonbirt 5.0456, USDA FS chi-16, JFK.02920, JFK.02908 and 2GD were hexaploid but JFK.02899 and USDA FS bar-2 (FRAX 28) were diploid. Kew’s 1992-377 and the samples in Gerry Douglas’ garden, all labelled as *F. mandshurica* but with anomalous barcode sequences, were hexaploid (unlike JFK 02932 identified by barcodes as *F. mandshurica* which had a diploid C-value). The other polyploids found were all samples for which the barcodes had failed to sequence: *F. biltmoreana* Westonbirt 5.0463 (FRAX17) was hexaploid, *F. profunda* Wakehurst 1993-1493 appeared to be octoploid, *F. lanuginosa* RBGE 20071296-G (FRAX22) was tetraploid, and *F. uhdei* USDA FS uhdei-ponto (FRAX34) was hexaploid; it is likely that the barcodes did not work well in these samples due to divergent homeologous copies of the barcode sequences among their sub-genomes.

### Fraxinus chiisanensis

Closer analysis of the morphology of the samples labelled as *F. mandshurica* at Kew and in Gerry Douglas’s garden, but with anomalous barcodes and hexaploidy, showed them to have naked buds which is characteristic of *F. chiisanensis* (Wallander, 2012), a rare endemic of Korea (Kim *et al*., 2022). The seeds for the trees in Gerry Douglas’ garden were sourced from the National Institute of Forest Science in Seoul,South Korea. A chromosome squash on a seedling from one of the Kew trees, which was hermaphroditic (1992-377), was also consistent with hexaploidy. A previous unpublished study also suggested that *F. chiisanensis* is hexaploid (mentioned in Siljak-Yakovlev *et al*., 2014).

Interestingly, the pattern of barcode polymorphism within these putative *F. chiisanensis* samples showed some unique alleles (OG9972: 1563, 1091, 1187, 764; OG8143: 390, 709) but also patterns of polymorphism involving alleles common in different sections (OG9972: 123, 153, 381, 405, 1048, 1487, 1698). In OG8143, *F. chiisanensis* shared alleles with *Melioides* at some loci (OG8143: 262-267, 801) but shared alleles with everything but *Melioides* at other loci (OG8143: 200, 237, 736-738). In OG22833, *F. chiisanensis* shared alleles with everything but section *Ornus* at 66-67, everything but *Melioides* at 144, everything but *Melioides* and *Dipetala* at 366, everything but *Ornus* and *Sciadanthus* at 372 and everything but *Pauciflorae* at 272. Thus, within its six genome copies, *F. chiisanensis* harbours a complex mix of variation found scattered across other sections of the genus.

### Hybrid crosses

Due to the barcoding and flow cytometry results reported above, several of the trees that were grafted for hybridisation were re-identified, including some that were used as parents in our crosses. *Fraxinus chinensis* (JFK.02906) was identified as diploid *F. ornus*. *Fraxinus japonica* (JFK.02920) (which, as noted above, is a synonym for *F. chinensis* (Wallander, 2012) was identified as the hexaploid *F. chinensis.* The trees labelled *F. mandshurica* in Dr Gerry Douglas’ garden were identified as hexaploid *F. chiisanensis*. *Fraxinus retusa henryana* (a synonym for *F. floribunda* in (Wallander, 2012))(JFK2005.0132), was confirmed as a diploid in section *Ornus* but could not be assigned to a species; we cannot be sure if it is *F. floribunda* or *F. ornus*. *Fraxinus pennsylvanica* (JFK.02931) was confirmed as being in section *Melioides* but could not be assigned to a species by the barcodes; we therefore have no reason to believe that the label is incorrect. *Fraxinus nigra* Marsh. × *F. mandshurica* (Canadian Hybrid No 8921 JFK.2001.0159) was not examined using barcodes, but we are confident that its parentage is reliably known (Davidson & Ronald, 2001).

Of the 35 crosses and controls that were performed (Supplementary Table 3), 756 seeds were produced, of which 363 were propagated, producing 131 seedlings of which 85 survived to be viable plants. We carried out an initial screening using the barcode region OG9927 on 84 seedlings that were alive at the time of the screening. This identified 16 putative hybrids showing heterozygosity at sites that varied among parental species. We then also sequenced these individuals with the barcoding regions OG22833 and OG8143. Region OG2833 was found to produce clear sequence reads of a comparable length to the average sequence length for the pure *Fraxinus* species. However, for the barcode region OG8143 we found only the forward primer was able to produce useable sequence reads for all of the putative hybrids, the useable region after sequence clean-up was found to be ∼650bp, this is ∼100bp short of the length of the full region.

None of the crosses that used non-native species as pollen donors produced hybrid offspring. In all of these cases, the offspring were sired by *F. excelsior* pollen. This includes all crosses involving *F. chiisanensis*, and flow cytometry confirmed that the progeny of the *F. excelsior* mother tree used in this cross were all diploid. There were 15 crosses done with non-native species as the maternal parent. Four of these produced no progeny. Seven of them had progeny that were confirmed as hybrid. We identified 14 intersectional hybrids; this included two plants of *F. pennsylvanica* □ *F. excelsior*, six of *F. ornus* □ *F. excelsior* and five section *Ornus* □ *F. excelsior.* For the latter five, the barcodes did not resolve whether one parent was *F. ornus* or *F. floribunda* within section *Ornus*.

Both of the *F. pennsylvanica* □ *F. excelsior* hybrids displayed vigorous growth and had no ash dieback symptoms in 2021, despite growing in an area with prevalent spore inoculum since 2018. One of these was still alive and healthy in July 2023. Most of the hybrids produced between *F. ornus* or section *Ornus* and *F. excelsior* died of ash dieback symptoms. Four were dead by the end of summer 2020. In 2021, one had severe symptoms, three had mild symptoms, and three were healthy. One was still alive and healthy in July 2023.

Sequence data for these two surviving hybrids for OG22833 and OG9927 are included in Supplementary Files 1 and 2. For OG9927, in the *F. pennsylvanica* □ *F. excelsior* hybrid, positions 416, 454, 603, 648, 743, 839, 1148, 1193 and 1482 were heterozygous. These sites are all diagnostic for section *Fraxinus* or section *Melioides* (see Table 2) confirming that this individual is a hybrid. In the section *Ornus* □ *F. excelsior* hybrid, positions 1148, 1465, 1482, 1538 and 1660 are heterozygous, confirming its hybridity between section *Ornus* and section *Fraxinus*. For OG22833, in the *F. pennsylvanica* □ *F. excelsior* hybrid, positions 144, 180 and 366 were heterozygous, which are diagnostic for *Melioides* (Table 2) and position 272. In the section *Ornus* □ *F. excelsior* hybrid, positions 66 and 67, which are diagnostic for section *Ornus* (Table 2), were heterozygous, as were positions 116, 266 and 332 which are fixed in one of the sections but polymorphic in the other.

### Spontaneous hybrids

We generated sequence data for four volunteer seedlings collected from the main ash area at Kew. Two of these were identified as *F. ornus* by at least one barcode (Supp. Table 2). Another was identified as being from Section *Melioides*. The fourth, which was a germinated seed lying in the grass when first collected, was identified as a hybrid between section *Fraxinus* and section *Melioides* by all three barcode regions and by OG8679. Two barcode regions identified *F. latifolia* as one parent. The seedling was collected close to *F. latifolia* 1981-8204 so this is likely to be a parent. The other parent was harder to identify but in barcode OG9927 it contained an allele found only in *F. excelsior* and *F. mandshurica*. In OG8679 it contained an allele found only in *F. excelsior*. Therefore we conclude that this individual is a hybrid between *F. latifolia* and *F. excelsior* (though we do not know the direction of the cross). It has survived its first summer outdoors in an ash dieback infested area without ash dieback damage but has not yet been fully challenged.

## Discussion

We have developed three low-copy nuclear gene barcodes that can delineate taxonomic sections and in some cases, individual species within the genus *Fraxinus*, allowing identification of ash samples and verification of hybrids. The cases where the barcodes did not perform well were certain polyploid species, and within large sections that contain many closely related species with a shared geographic distribution. For identification of the latter, use of whole genome sequencing (Kelly *et al*., 2020; Liu *et al*., 2024), AFLPs (Hinsinger *et al*., 2014) or SNP arrays may help, but the large amount of gene flow that occurs among some species of *Fraxinus* may still mean that diagnostic alleles are rare. Knowledge of genome-wide variation within many *Fraxinus* species is currently limited, but where it has been surveyed in the case of *F. excelsior, F. angustifolia and F. mandshurica*, it has shown widespread sharing of alleles and even raised questions as to whether they are separate species (Hinsinger *et al*., 2014). Thus, the difficulty we have in separating species in the section *Fraxinus* with our barcodes is a biological result, not a shortcoming of the barcodes. Barcoding of larger sample sizes in the future could identify more alleles found exclusively in a subset of populations within some species. Sample identifications using the barcodes sometimes differed from living collection labels but seldom differed from identifications by Eva Wallander.

Use of these barcodes on living collections suggests that mislabelling is not uncommon in living collections. Potential mis-labelling of Asiatic species may reflect lower acquaintance of European curators with Asiatic species than with European and American species. Several cases of mislabelling in living collections involve a local species being given an exotic label: in some cases this may be because exotic scion material had been grafted to a local species rootstock, but shoots from the rootstock have become dominant and the exotic material has not survived. Other cases of mislabelling may be *ad hoc* errors within collections. Our findings highlight the need for verification of living collection labels before undertaking experiments or initiating breeding programmes. The rare Korean endemic *F. chiisanensis* is commonly labelled as *F. mandshurica*, though a seedlot of *F. mandshurica* supplied by Sandeman Seeds (Dalkey, Ireland) was found to be correctly labelled. Samples we identify as *F. mandshurica* are sometimes labelled as *F. platypoda*, but there seems to be genuine ambiguity around the latter species, as Kim et al (2022) found Chinese samples of *F. platypoda* to cluster among *F. mandshurica* samples (see also Jeandroz *et al*., 1997), whereas Japanese samples of *F. platypoda* were in a clade sister to *F. chiisanensis*.

We generated and verified wide hybrids of ash, among sections *Fraxinus*, *Melioides* and *Ornus*: these were from *F. pennsylvanica* □ *F. excelsior* and section *Ornus* □ *F. excelsior* crosses. We also discovered a spontaneous hybrid formed between *F. latifolia* (section *Melioides*) and *F. excelsior* (section *Fraxinus*) at Kew Gardens. Many hybrids have been reported or made previously within the genus *Fraxinus* (see Introduction) but ours is the first study we are aware of to have confirmed the production of intersectional hybrid plants between sections *Ornus* and *Fraxinus*. Unlike the wide hybridisations recently conducted between sections *Fraxinus* and *Melioides* (*F. mandshurica* □ *F. americana*, and *F. mandshurica* □ *F. velutina*)(Zeng *et al*., 2015; He *et al*., 2019), we did not use electrostatic treatments to overcome fertility barriers.

One of our major objectives when setting up crosses was to hybridise *F. excelsior* with east Asian ash species possessing high resistance to both EAB and ADB. We have not achieved this, partly due to initial misidentification of parental trees, some of which were labelled as Asiatic species, but turned out not to be; especially *F. chiisanensis* which was believed to be *F. mandshurica* at the time of crossings. We attempted many crosses with *F. chiisanensis* pollen, but these were unsuccessful. We might have had more success by using *F. excelsior* as a pollen donor, since hybridisation and introgression can be asymmetric, as in oaks (Petit *et al*., 2004); however no female trees of *F. chiisanensis* were available. We cannot exclude the possibility that there are intrinsic incompatibilities between *F. excelsior* and *F*.

*chiisanensis.* While *F. excelsior* is a diploid, *F. chiisanensis* is hexaploid and our barcodes suggest it could have a complex hybrid origin.

Whilst the majority of our hybrids died from ash dieback between 2018 and 2023, one individual from each of our crosses remained alive and healthy in summer 2024. Although *F. pennsylvanica* is highly susceptible to EAB (Koch *et al*., 2015; Orlova-Bienkowskaja & Bieńkowski, 2016), it does appear to be less susceptible to ADB than *F. excelsior* (Nielsen *et al*., 2016; Drenkhan *et al*., 2017; Plumb *et al*., 2019). Thus, crosses between *F. pennsylvanica* and *F. excelsior* may have promise for ADB resistance breeding, especially as *F. pennsylvanica* is a good timber species. Section *Ornus* contains species with low susceptibility to ADB and EAB (Nielsen *et al*., 2016; Drenkhan *et al*., 2017; Plumb *et al*., 2019; Kelly *et al*., 2020), so hybrids from this section may also be of interest. The spontaneous hybrid between *F. latifolia* and *F. excelsior* has survived its first summer outdoors in an ash dieback infested area without incurring ash dieback damage. Although *F. latifolia* is highly susceptible to EAB (Kelly *et al*., 2020), it appears to be less susceptible to ADB than *F. excelsior* (Nielsen *et al*., 2016; Drenkhan *et al*., 2017; Plumb *et al*., 2019), so the hybrid may have some promise.

Spontaneous hybrids in living collections are often seen as a liability and a concern for *ex situ* conservation (M Maunder, C Hughes, JA Hawkins, A Culham, 2004; Ensslin & Godefroid, 2019). However, with means to readily identify them and their parentage, they could be a valuable resource. Their discovery may save much effort in setting up artificial pollinations and breaking seed dormancy. Here, in a sample of just four volunteer seedlings from Kew Gardens, we identified an intersectional hybrid between the American species *F. latifolia*, and the European species *F. excelsior*. Wider sampling of seeds and seedlings in botanic gardens and arboreta may yield many more hybrids.

The barcodes and hybrids presented here provide methodological and biological resources for the use of world-wide ash germplasm for resistance to pests and pathogens and adaptation to climate change. The success of wide hybridisations suggests that mobilisation of genetic diversity throughout the genus in hybrid breeding programmes may be possible for the genus *Fraxinus*. The development of hybrid breeding programmes between *F. excelsior* and non-native species, may be part of the solution to the threats of ADB and EAB. In addition, the more extensive field testing of confirmed ash species (using barcodes) from diverse regions of provenance may identify adapted material for commercial timber production and rapid carbon sequestration.

## Acknowledgements

We thank Eva Wallander for her expert identifications of numerous samples used in this study. We thank Jaume Pellicer for advice and assistance with flow cytometry. We thank Timothy Baxter, Samuel Brockington, Peter Brownless, Dan Crowley, Dawn Edwards, Simon Honey, Ross Irvine, Richard Jinks, Penny Jones, Simon Toomer, Tony Kirkham, Hugh McAllister, Iain Parkinson and Sara Redstone for help with obtaining *Fraxinus* materials from UK collections. We thank Colin Kelleher, National Botanic Gardens of Ireland, Glasnevin for provision of scion wood of exotic species and John McNamara for technical assistance in grafting. R.J.A.B. and L.J.K. acknowledge funding from the Erica Waltraud Albrecht Endowment Fund and the Living with Environmental Change Tree Health and Plant Biosecurity Initiative – Phase 2 (grant no. BB/L012162/1), funded jointly by BBSRC, Defra, ESRC, Forestry Commission, NERC and the Scottish Government. W.J.P. was supported by a Walsh Fellowship from Teagasc, and funding from Defra.

## Author contributions

R.J.A.B., W.J.P., L.J.K, J.K. and G.C.D. planned and designed the research. W.J.P., J.M., D.C., L.C. and W.C. extracted DNA. W.J.P and L.C. conducted Sanger sequencing. R.F.P. conducted flow cytometry. G.C.D. conducted hybridisations. W.J.P., L.J.K, M.N.G. and J.K. collated samples.

R.J.A.B., W.J.P., L.J.K, J.K. and G.C.D. wrote the manuscript.

## Supplements

**Supplementary Table 1.** Table of barcoded samples and their results (Excel format)

**Supplementary Table 2.** Table of hybrid crosses and their results (Excel format)

**Supplementary File 1.** Multi-sequence alignment for OG9972 (fasta format)

**Supplementary File 2.** Multi-sequence alignment for OG22833 (fasta format)

**Supplementary File 3.** Multi-sequence alignment for OG8143 (fasta format)

**Supplementary File 4.** Multi-sequence alignment for OG8679 (fasta format)

